# Enhanced mismatch negativity in harmonic compared to inharmonic sounds

**DOI:** 10.1101/2021.10.15.464562

**Authors:** D. R. Quiroga-Martinez, K. Basiński, J. Nasielski, B. Tillmann, E. Brattico, F. Cholvy, L. Fornoni, P. Vuust, A Caclin

**Author notes:** Equally contributing authors (P. Vuust, A Caclin).

## Abstract

Many natural sounds have frequency spectra composed of integer multiples of a fundamental frequency. This property, known as harmonicity, plays an important role in auditory information processing. However, the extent to which harmonicity influences the processing of sound features beyond pitch is still unclear. This is interesting because harmonic sounds have lower information entropy than inharmonic sounds. According to predictive processing accounts of perception, this property could produce more salient neural responses due to the brain’s weighting of sensory signals according to their uncertainty. In the present study, we used electroencephalography to investigate brain responses to harmonic and inharmonic sounds commonly occurring in music: piano tones and hi-hat cymbal sounds. In a multi-feature oddball paradigm, we measured mismatch negativity (MMN) and P3a responses to timbre, intensity, and location deviants in listeners with and without congenital amusia—an impairment of pitch processing. As hypothesized, we observed larger amplitudes and earlier latencies (for both MMN and P3a) in harmonic compared to inharmonic sounds. These harmonicity effects were modulated by sound feature. Moreover, the difference in P3a latency between harmonic and inharmonic sounds was larger for controls than amusics. We propose an explanation of these results based on predictive coding and discuss the relationship between harmonicity, information entropy, and precision weighting of prediction errors.

## Introduction

Many naturally occurring sounds are comprised of frequencies that are integer multiples of a single, fundamental frequency (F0). These are termed *harmonic* sounds and include human and animal voices as well as many musical instrument sounds. Harmonic sounds have an easily detectable pitch, a subjective perceptual quality that enables them to be ordered from low to high (McDermott, 2018; Oxenham, 2012). Pitch information is used for solving many cognitive tasks. In tonal languages, pitch contours determine the meaning of words, while in non-tonal languages they are used for prosody. In music, pitch is critical for the processing of melody and harmony (Huron, 2016). Pitch information is also used in auditory scene analysis, where simultaneously perceived sounds can be segregated using pitch discrepancies (Bregman, 1994; Wightman & Green, 1974). *Inharmonic* sounds possess frequency spectra that do not follow the harmonic series. These are often described as metallic, crackling, sizzling or noisy. Inharmonic sounds are sparingly used in Western tonal music (mainly in percussion instruments) as they do not produce a clear pitch sensation that may be used to convey melody.

Historically, research in auditory neuroscience focused on simple stimuli, mostly limited to sine waves and harmonic complex tones (McPherson & McDermott, 2020a). Not much is known about the neural mechanisms of perception of inharmonic sounds or how inharmonicity influences the perception of auditory features other than pitch. Recent studies have shown harmonicity to be an important factor in several auditory tasks not directly related to pitch. Violations of harmonicity impair speech recognition in cocktail-party scenarios (Popham, 2018). Harmonic sounds are easier to discriminate than inharmonic sounds over time delays, suggesting more efficient coding of harmonic sounds in memory (McPherson & McDermott, 2020b). Harmonic signals are also more easily detected in noise than are inharmonic signals, pointing to an important role of harmonicity in auditory scene analysis (McPherson et al., 2020).

Taken together, these studies suggest that harmonicity is important for processing different auditory features. It is thus likely that the ease of detection of deviances in features, such as timbre, intensity or location, could depend on harmonicity. This view is also consistent with a predictive coding account of auditory perception, where sensory information, usually cast as ascending prediction error, is weighted by the information entropy (i.e., uncertainty or inverse precision) of the stimuli (Clark, 2015; Friston, 2005; Rao & Ballard, 1999). As inharmonic sounds have higher spectral entropy, prediction errors elicited by inharmonic deviants may be down-weighted in comparison to harmonic deviants.

In this study, we used electroencephalography to investigate the brain responses to harmonic and inharmonic sounds in both typical listeners and participants with congenital amusia (CA) — an impairment in pitch processing that cannot be attributed to causes such as intellectual disability, lack of music exposure, or brain damage (Peretz, 2016).

Aiming for ecological validity, our present study used sounds commonly occurring in popular music: piano tone (harmonic, low spectral entropy) and hi-hat cymbal (inharmonic, high spectral entropy). In a multi-feature oddball paradigm (Näätänen et al., 2004; Paavilainen, 2013), we measured mismatch negativity (MMN) and P3a responses to timbre, intensity, and location deviants. These ERPs are elicited without attentional control and index the brain’s ability to perform automatic comparisons of incoming auditory stimuli (Garrido et al., 2009; Paavilainen, 2013). In contrast to traditional paradigms that employ a single type of deviant, multi-feature paradigms record MMN responses to several sound features in a relatively short time, which make them an optimal choice to address our question (Kliuchko et al., 2016; Vuust et al., 2011). Based on previous studies highlighting the role of harmonicity in auditory scene analysis, we hypothesized that timbre, intensity, and location deviants would elicit larger MMN and P3a responses for harmonic than for inharmonic sounds. Finally, given that listeners with congenital amusia are impared in the processing of pitch information (which relies on harmonicity), here we also explored whether amusics process harmonic and inharmonic sounds differently from their matched non-musician controls (Cousineau et al., 2012; Marin et al., 2012).

## Methods

The code and materials employed to conduct the experiment and analyses presented here can be found at: https://doi.org/10.17605/OSF.IO/JSEU8. Due to data protection regulations, data cannot be publicly shared, but can be made privately available upon reasonable request.

### Participants

Thirty-four participants took part in the experiment (same participants as in Quiroga-Martinez et al., 2021), 17 amusics and 17 matched controls (see Table 1). All subjects were recruited in the Lyon area in France, were French speakers, and were screened for congenital amusia using the Montreal Battery of Evaluation of Amusia (MBEA) (Peretz et al., 2003). The total MBEA scores including six subtests were significantly lower for amusics than for controls, t(25.6) = −10.49, p < .001; as were the average combined scores for the three pitch subtests, t(23.7) = −11.36, p < .001. An individual was considered amusic if the total MBEA score was less than 23 (maximum score = 30) or their average score for the pitch subtests was lower than 21.7 (maximum score = 30). We measured pitch discrimination thresholds (PDT) using an adaptive tracking staircase procedure (Tillmann et al., 2009). PDTs were significantly larger for amusics in comparison to controls. The groups did not differ significantly in age, years of education, or musical training. The study was conducted and approved by a national ethics committee. Participants gave their written informed consent prior to the experiment and received a small financial compensation for their time.

**Table 1.** Participant demographics (mean ± SD, *t* statistics, degrees of freedom and *p*-values). MBEA: Montreal Battery of Evaluation of Amusia (maximum score= 30, average of the six sub-tests of the battery); MBEA pitch: average of the three sub-tests of the battery assessing pitch (maximum score= 30); PDT: Pitch Discrimination Threshold. For music training, Mann-Whitney U test results are reported.

|  | **Amusics** | **Controls** | ***t*** | ***df*** | ***p*** |
| --- | --- | --- | --- | --- | --- |
| Sample size | 17 | 17 | - | - | - |
| Female | 8 | 9 | - | - | - |
| Right-handed | 13 | 14 | - | - | - |
| Education (years) | 15.06 (± 2.7) | 15.12 (± 2.34) | -0.07 | 31.4 | .946 |
| Music training (years) | 0 | 0.24 (± 0.66) | 127.5 (U) | - | .163 |
| Age (years) | 38.43 (± 15.96) | 37.61 (± 17.02) | 0.15 | 31.9 | .886 |
| MBEA | 21.79 (± 1.81) | 27.09 (± 1.04) | -10.49 | 25.6 | <.001 |
| MBEA pitch | 20.79 (± 2.15) | 27.43 (± 1.09) | -11.36 | 23.7 | <.001 |
| PDT (semitones) | 1.57 (± 1.53) | 0.31 (± 0.17) | 3.375 | 16.4 | .004 |

### Stimuli

Two types of sound sequences were presented to participants, corresponding to a modified version of the optimal MMN paradigm (Näätänen et al., 2004). In the piano condition, participants listened to repeating piano tones with a fundamental frequency at C3 pitch (F0 = 262 Hz). In the hi-hat condition, broad-band percussive hi-hat cymbal sounds were used (Figure 1). The sounds were samples taken from the Cubase sample library (Steinberg Media Technology, version 8). The spectral entropies of the sounds were 2.94 bits and 9.66 bits for the piano and hi-hat tones, respectively (as calculated with AntroPy v.0.1.4; Vallat, 2021). During the experiment, one sound was played repeatedly, and a deviant was pseudorandomly introduced in every group of four consecutive stimuli in the sequence. No two deviants were played consecutively, and no deviant feature was presented again before a whole iteration of the five features was played. Each condition (piano or hihat) was presented in a separate block lasting approximately 13 minutes. Both sequences included the same number of standard and deviant tones. The piano block and the hi-hat block, together with two other conditions involving melodies with different complexity levels (presented in Quiroga-Martinez et al., 2021), were counterbalanced between participants and their order matched across groups. Other conditions were included in the experiment, which have been reported elsewhere (Quiroga-Martinez et al., 2021).

**Figure 1.**
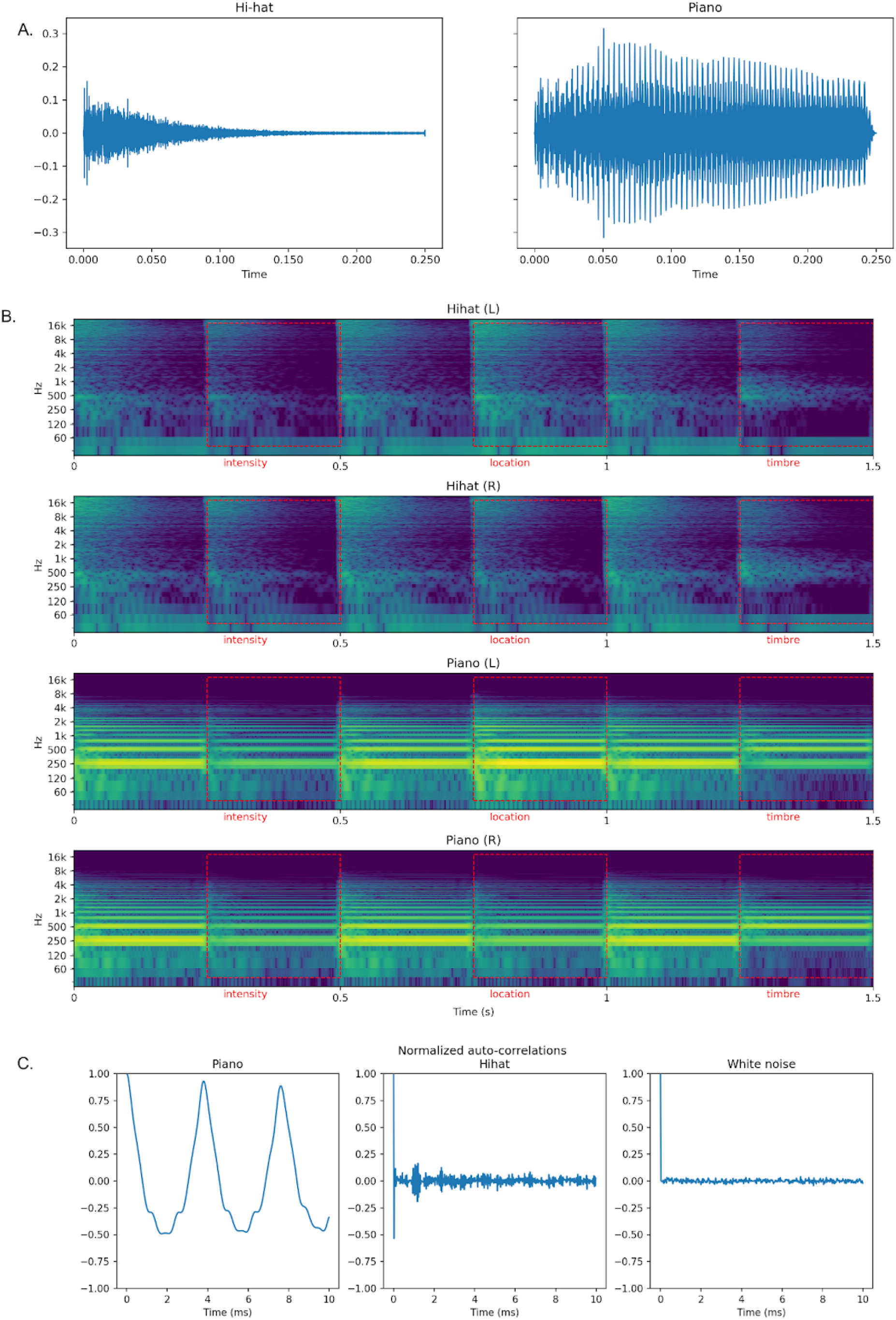
(A) Waveforms for standard hi-hat and piano sounds. (B) Example of the stimulus deviants in each condition for each feature (red rectangles), represented as a power spectrogram showing left and right audio channels for two sound conditions. Vertical axes show (log-spaced) frequency while horizontal axes show the time in seconds. For every condition, standard sounds are intertwined with different feature deviants. Note: the order of the stimuli in this figure is for illustration purposes, see main text for stimulus ordering procedures. (C) Normalized auto-correlations of standard piano and hi-hat sounds in comparison with white noise. Piano sounds are much more periodic than hi-hat sounds and white noise.

The sounds lasted 250 ms each, were loudness normalized, and were presented excluding any silent gaps between the sounds. Pitch, intensity, timbre, location, and rhythm deviants were introduced in the melodies. Pitch deviants appeared in the piano condition but were not part of the analysis because there are no pitch deviants in the hi-hat condition (results for the pitch deviant in the piano block can be seen in Figure 2 in Quiroga-Martinez et al., 2021). All deviants were created with Adobe Audition (Adobe Systems Incorporated, version 8) by modifying the standard tones as follows: intensity: −12 dB gain; timbre: a combination of filters (low-shelf −10 dB at 500 Hz, peak +10 dB at 2 kHz, notch filter at 6 kHz, all filters Q = 1); location: leftward bias (20 ms time shift between channels); rhythm: −60 ms for sound onset. Note that rhythm violations implied a shortening of the preceding tone and a lengthening of the actual deviant tone by 60 ms. The rhythm deviants were excluded from the analysis due to baseline contamination issues, as reported in Quiroga-Martinez et al. (2021). Overall, a total of 2339 standards and 153 (5%) deviants per feature were presented in each of the two conditions.

**Figure 2.**
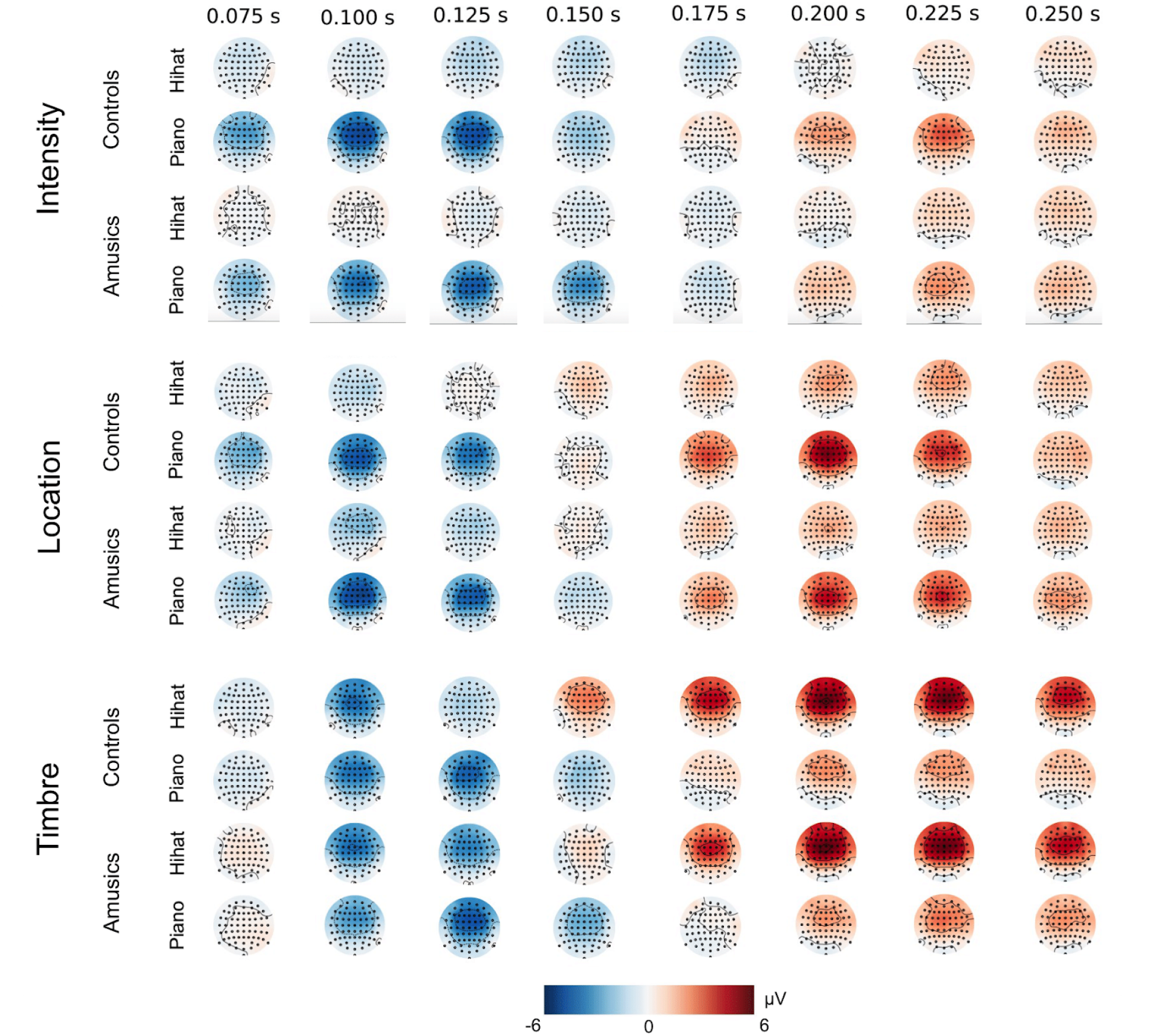
Time-course of grand-average MMN (blue) and P3a (red) topographies for different features (location, timbre, and intensity), groups (amusics and controls) and conditions (piano and hi-hat)

### Procedure

Participants were informed about the procedures at their arrival, willingly gave their written consent and filled out the required forms. Subsequently, EEG caps were placed on their scalp accompanied by conductive gel applied to the electrodes. A Sennheiser HD280 Pro headset was carefully placed on top of the EEG cap with foam padding to avoid pressure on electrodes. The impedances were checked again after the headphones were on. All participants had the sound volume set to an identical, comfortable level. During testing, participants were looking at a computer screen from a distance of about 1.5 m, sitting on an armchair inside a sound-attenuated booth, electrically-shielded with a Faraday cage. They were told sounds would be playing in the background and were instructed to watch a movie of their choice while ignoring the auditory stimuli. Additionally, they were asked to remain still and relaxed, knowing there would be pauses between blocks during which they could stretch and change posture. During stimulation, the blocks were presented in such a way that for 9 matched pairs of participants, the counterbalanced piano and hi-hat conditions, together with the two additional complexity conditions, came before two counterbalanced additional conditions, whereas for the remaining pairs the order was inverted. Two additional blocks were included at the end of the experiment, in which participants listened freely to entire pieces of music. Their analysis, however, is beyond the scope of this article and will be reported elsewhere. The whole recording session lasted around one hour and a half, plus half an hour of preparation.

### EEG recording and preprocessing

Scalp potentials were recorded with a 64-channel Biosemi system with active electrodes and a sampling rate of 1024 Hz. Additional electrodes were used to track horizontal and vertical eye movements. Data analyses were conducted with MNE-Python v.0.23.0 (Gramfort et al., 2014). EEG signals were first cleaned from eyeblink artifacts using independent component analysis with a semiautomatic routine (fastICA algorithm). Visual inspection was used as a quality check. After removing ICA components, the raw signals were filtered with a pass band of 0.5 - 35 Hz and re-referenced to the mastoids. Epochs from −100 ms to 400 ms from tone onsets were extracted and baseline corrected with a prestimulus baseline of 100 ms. Epochs with an amplitude exceeding 150 μV were rejected to further clean the data from remaining artifacts. For each participant, event-related potentials (ERP) were obtained by averaging epochs for the standard tones and each of the deviant features separately, per condition. Standard tones preceded by a deviant were excluded from the averages. Both deviant-specific MMN and P3a responses were calculated by subtracting standard from deviant ERPs for each feature and condition.

### Statistical analyses

We performed analyses on mean amplitudes and latencies for both MMN and P3a responses. MMN peak latencies were extracted within a time window of 70 - 250 ms, whereas P3a peak latencies were extracted for 150 - 350 ms. Mean amplitudes were obtained from electrodes Fz, F1, F2, F3, F4, FCz, FC1, FC2, FC3, FC4 and calculated as the average activity ± 25 ms around the participant-wise peak, for each condition (piano or hi-hat) and feature (intensity, location, timbre). The chosen electrodes correspond to those that typically show the largest responses in MMN studies and exhibited the largest P50 amplitudes (thus making sure that they properly captured auditory evoked activity). Using R v. 4.0.5 (R Core Team, 2021) and the ezANOVA library (Lawrence, 2016), several mixed ANOVAs including within- and between-subject effects were run on the mean amplitudes and latencies of MMN and P3a difference waves. Factors included group (amusics and control), condition (piano and hi-hat), and sound feature (intensity, location, and timbre). Mauchly’s test of sphericity was checked for each ANOVA and no correction was applied. Post-hoc, pairwise contrasts were also performed with the emmeans library (Lenth et al., 2021) after fitting linear mixed-effects models using lme4 library (Bates et al., 2015) with the two main effects and their interactions as predictors.

## Results

Consistent with the literature (Näätänen et al., 2007, Polich & Criado, 2006), the MMN and P3a manifested themselves as a fronto-central negativity and a fronto-central positivity, respectively (Figure 2 and 3).

**Figure 3.**
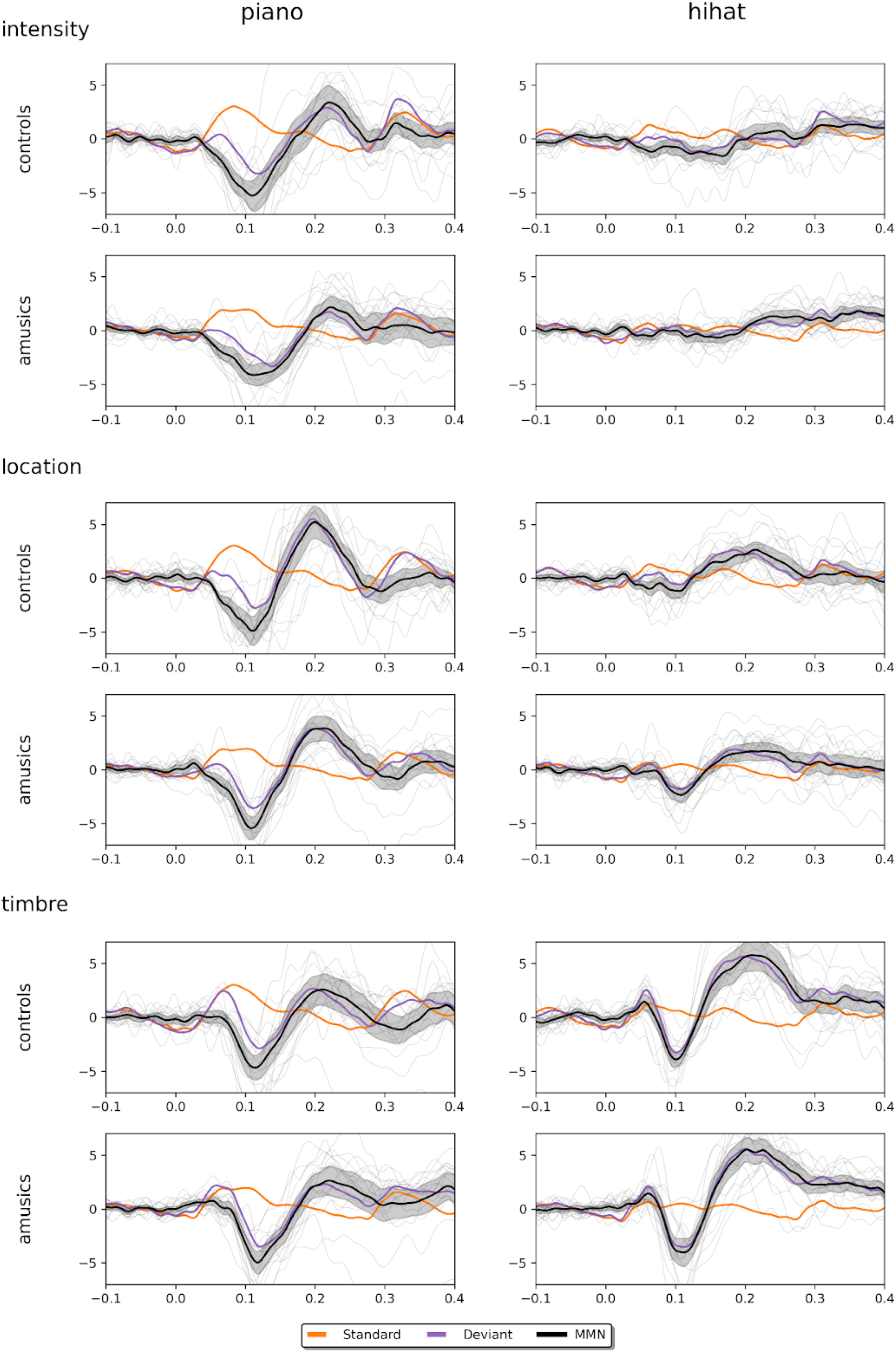
Standard, deviant and MMN difference waves, averaged according to conditions, groups and features. Vertical axes show neural activity in microvolts and horizontal axes show the time in seconds. Gray traces represent MMN difference waves for each participant. Shaded areas indicate 95% confidence intervals (for the difference waves). The displayed activity corresponds to the channel Fz.

**Figure 4.**
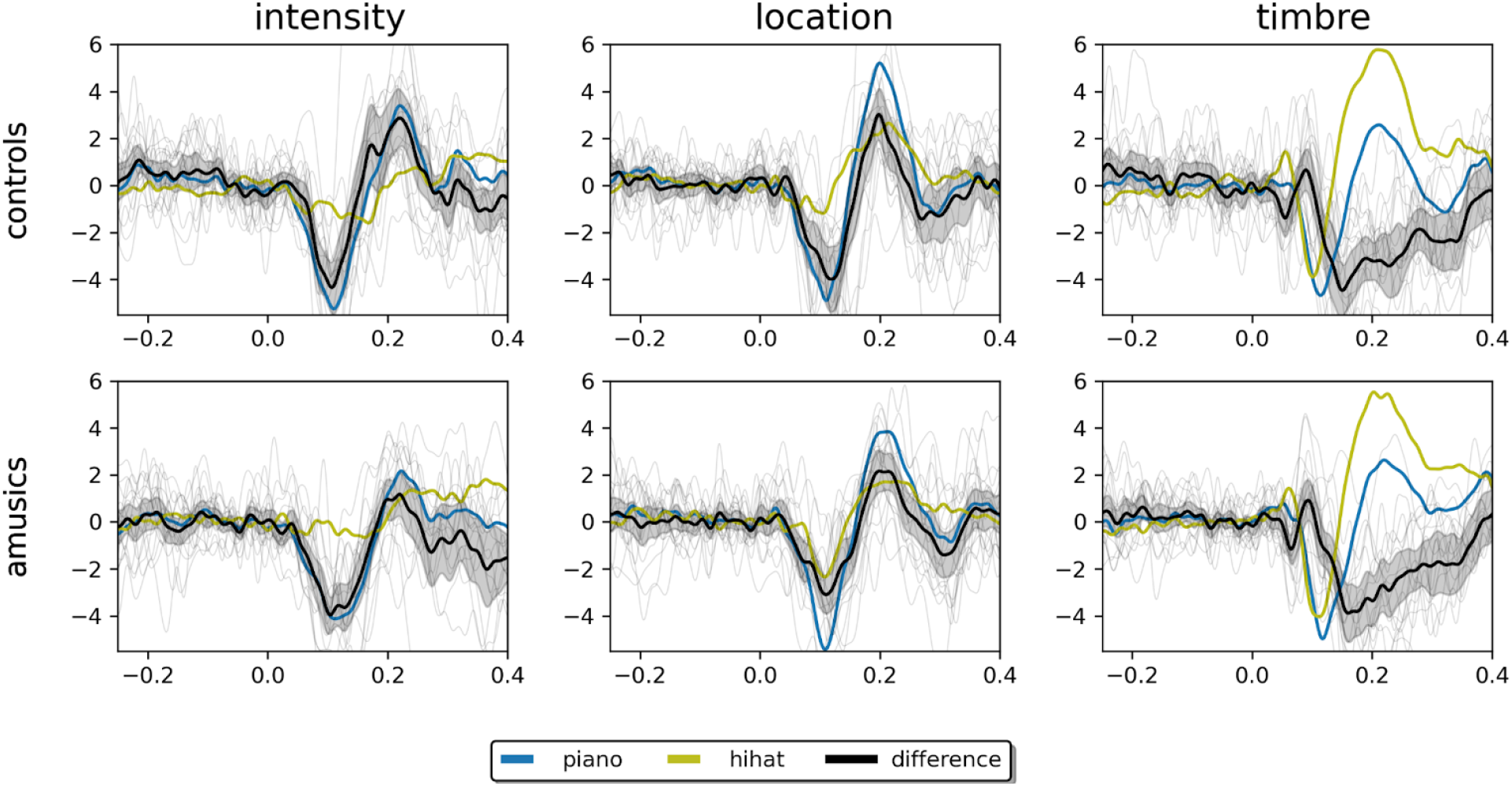
Effect of condition on MMN and P3a responses for each group. Gray traces represent difference waves between piano and hi-hat condition for each participant. Shaded areas indicate 95% confidence intervals (for the difference of difference waves). The displayed activity corresponds to the channel Fz.

### MMN analysis

Mixed effects ANOVA analyses revealed significant main effects and two-way interactions on MMN mean amplitudes and peak latencies for group, condition, and feature (Figure 5).

**Figure 5.**
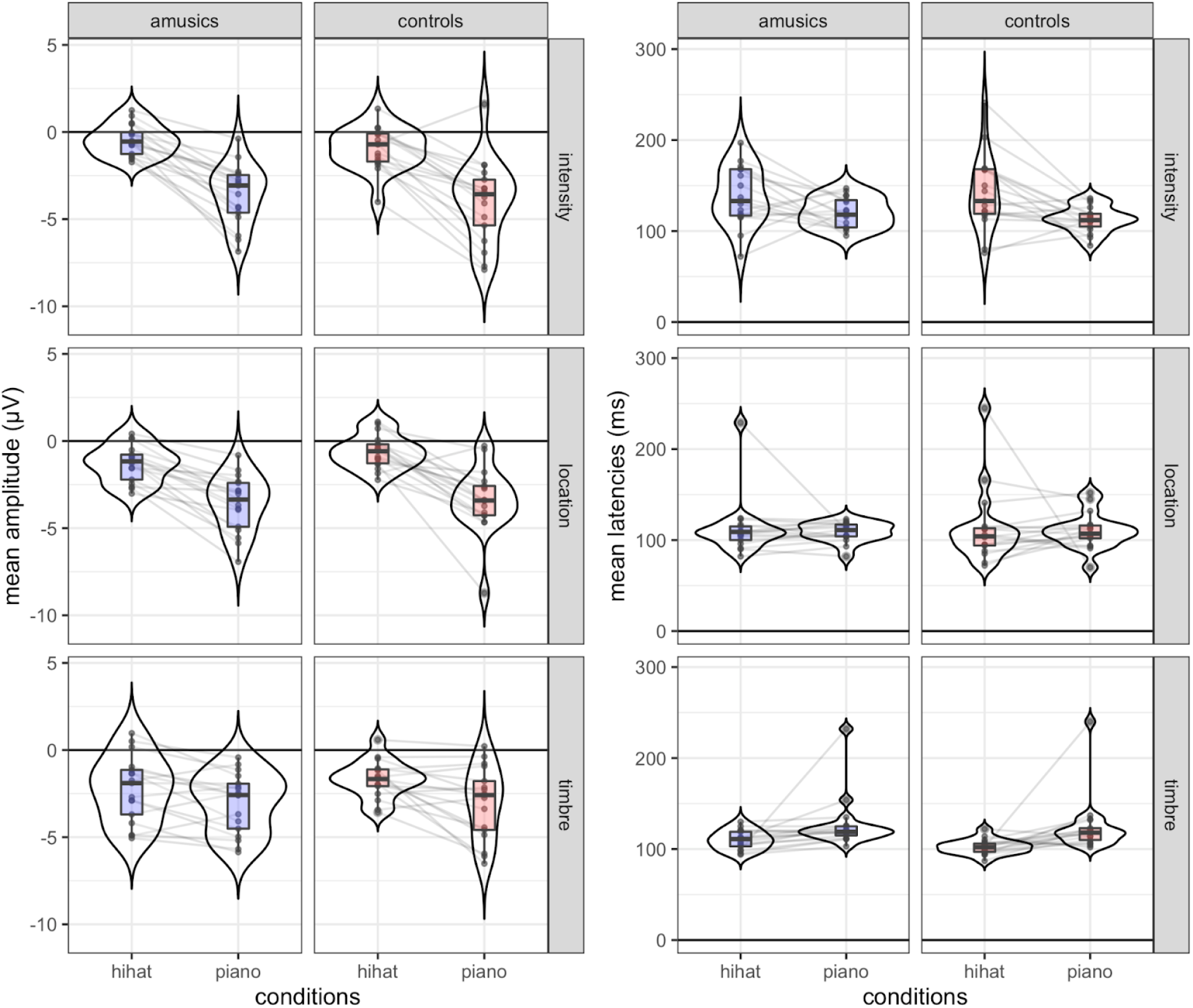
Mean MMN amplitudes (left) and peak latencies (right) as a function of conditions and features in both groups. Boxes display median and interquartile ranges. Beans depict the estimated densities. Lines connect measurements for individual participants.

For mean amplitudes (see Appendix 1), the main effect of condition was significant F(1,32) = 101.129, η^2^ = 0.338, p < .001, with larger MMNs for piano tones compared to hi-hat tones. No significant main effects of group and feature were found, with F(1,32) = 0.063, p = .803 and F(2,64) = 1.131, p = .329 respectively. No significant interaction was found between group and condition F(1,32) = 0.486, p = .491. Furthermore, the interaction between group and feature was not significant either, with F(2,64) = 2.561, p = .085. Interestingly, a significant interaction was found for condition and feature with F(2,64) = 24.485, η^2^ = 0.068, p < .001. No significant three-way interaction between group, condition and feature was observed, F(2,64) = 0.775, p = .465.

By looking at the post hoc pairwise comparison (Table 2), we can observe that there are significant differences in amplitude, contrasting hi-hat and piano conditions, for all features. We notice in Table 2 that the estimates of between-condition MMN differences (hi-hat - piano) for intensity and location were relatively high compared to timbre. This is reflected in the significant differences in between-condition contrasts comparing intensity to timbre (p < 0.001) and location to timbre (p = 0.001) but not intensity to location (p = 0.504). Taken together, these differences account for the significant condition-by-feature interaction.

**Table 2.** Pairwise contrasts of MMN mean amplitudes between conditions for each feature separately, as well as pairwise comparisons between features for differences between conditions (contrasts of contrasts). Data is pooled from both groups. Significant contrasts are highlighted in bold and marked with an asterisk. Standard effect sizes (d) are calculated as the difference between conditions divided by the square root of the sum of the residual and the random effects variance. The condition by feature interaction was not further modulated by group, p = .465.

| Feature | Contrast | Difference ($\mu V$) | CI 2.5% | CI 97.5% | t | d | p |
| --- | --- | --- | --- | --- | --- | --- | --- |
| Intensity | hi-hat - piano | 3.10 | 2.53 | 3.66 | 10.82 | 1.92 | **<0.001\*** |
| Location | hi-hat - piano | 2.54 | 1.97 | 3.10 | 8.87 | 1.57 | **<0.001\*** |
| Timbre | hi-hat - piano | 1.09 | 0.52 | 1.65 | 3.81 | 0.67 | **<0.001\*** |
| (hi-hat - piano intensity) -<br>(hi-hat - piano location) |  | 0.56 | -0.42 | 1.54 | 1.39 | 0.35 | 0.504 |
| (hi-hat - piano intensity) -<br>(hi-hat - piano timbre) |  | 2.01 | 1.03 | 2.99 | 4.96 | 1.24 | **<0.001\*** |
| (hi-hat - piano location) -<br>(hi-hat - piano timbre) |  | 1.45 | 0.47 | 2.43 | 3.58 | 0.9 | **0.001\*** |

For mean latencies (see Appendix 2), the main effect of feature was significant F(2,64) = 9.252, η^2^ = 0.073, p < .001. No significant main effects were found for group and condition, with F(1,32) = 0.195, p = .662 and F(1,32) = 0.475, p = .496 respectively. No significant interaction was found for group-by-condition F(1,32) = 0.064, p = .802 or group-by-feature F(2,64) = 0.252, p = .778. Interestingly, an interaction was found for condition and feature with F(2,64) = 11.520, η^2^ = 0.105, p < .001. No significant three-way interaction between group, condition and feature was observed, F(2,64) = 0.457, p = .635.

By looking at the post-hoc pairwise comparisons decomposing the condition-by-feature interaction (Table 3), we can observe that there are significant differences in latencies, contrasting hi-hat and piano conditions, for the intensity with 23.62 milliseconds (earlier MMN for piano tones, p < 0.001) and timbre with −19.68 milliseconds (earlier MMN for hi-hat tones, p = 0.002). No significant differences in latency between conditions were found for location (p = 0.570). Similarly to MMN mean amplitudes (Table 2), the condition-by-feature interaction could be explained with significant differences in between-condition contrasts comparing intensity to timbre (p < 0.001) and location to timbre (p = 0.025) but not intensity to location (p = 0.065).

**Table 3.** Pairwise contrasts of MMN peak latencies between conditions for each feature separately, as well as pairwise comparisons between features for differences between conditions (contrasts of contrasts). Data is pooled from both groups. Significant contrasts are highlighted in bold and marked with an asterisk. Standard effect sizes (d) are calculated as the difference between conditions divided by the square root of the sum of the residual and the random effects variance.

| Feature | Contrast | Difference (ms) | CI 2.5% | CI 97.5% | t | d | p |
| --- | --- | --- | --- | --- | --- | --- | --- |
| Intensity | hi-hat - piano | 23.62 | 11.49 | 35.75 | 3.85 | 0.89 | **<0.001\*** |
| Location | hi-hat - piano | 3.50 | -8.63 | 15.63 | 0.57 | 0.13 | 0.570 |
| Timbre | hi-hat - piano | -19.68 | -31.81 | -7.55 | -3.20 | -0.74 | **0.002\*** |
| (hi-hat - piano intensity) -<br>(hi-hat - piano location) |  | 20.12 | -0.9 | 41.13 | 2.32 | 0.76 | 0.065 |
| (hi-hat - piano intensity) -<br>(hi-hat - piano timbre) |  | 43.29 | 22.28 | 64.31 | 4.98 | 1.63 | **<0.001\*** |
| (hi-hat - piano location) -<br>(hi-hat - piano timbre) |  | 23.18 | 2.16 | 44.19 | 2.67 | 0.87 | **0.025\*** |

### P3a analysis

Mixed ANOVA analyses revealed significant main effects and two-way interactions on P3a mean amplitudes and peak latencies for group, condition, and feature (Figure 6).

**Figure 6.**
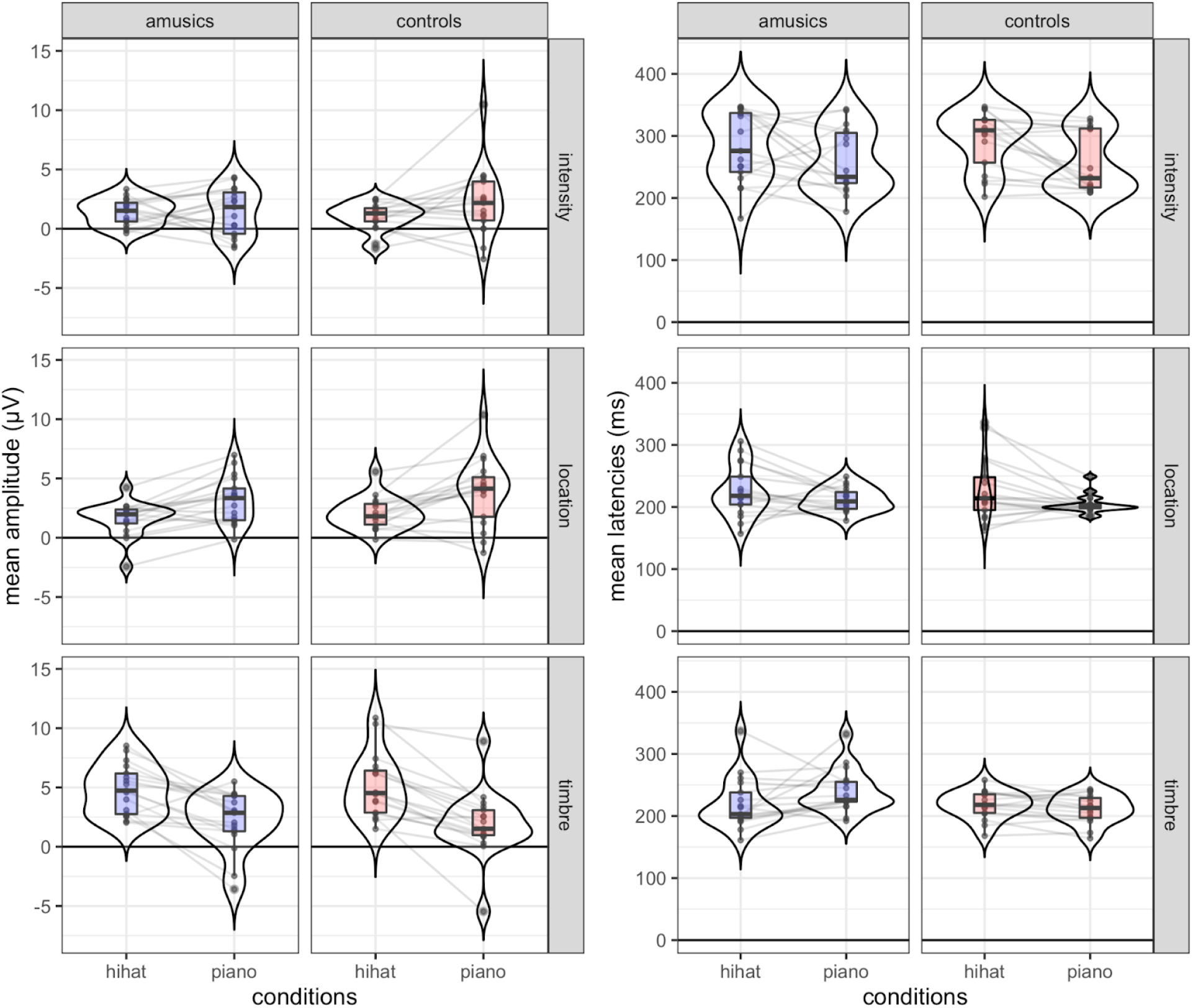
Mean P3a amplitudes (left) and peak latencies (right) as a function of conditions and features in both groups. Boxes display median and interquartile ranges. Beans depict the estimated densities. Lines connect measurements for individual participants.

For P3a mean amplitudes (see Appendix 3), the main effect of feature was significant with F(2,64) = 24.580, η^2^ = 0.130, p < .001. No significant main effects of group and condition were found F(1,32) = 0.223, p = .640 and F(1,32) = 0.863, p = .360. Interestingly, a significant condition-by-feature interaction was found with F(2,64) = 44.171, η^2^ = 0.180, p < .001. No significant interactions were found for group-by-condition with F(1,32) = 0.341, p = .563 and group-by-feature with F(2,64) = 0.518, p = .598. No significant three-way interaction was found for group-by-condition-by-feature with F(2,64) = 1.493, p = .232.

By looking at the post-hoc pairwise comparisons (Table 4), we can observe that there is a significant difference in amplitude, both for location and timbre in both groups, contrasting hi-hat and piano conditions. However, whereas for location, P3a was smaller for hi-hat tones compared to piano tones (as already observed for the MMN), for timbre P3a was larger for hi-hat tones compared to piano tones. This is reflected in the significant difference between location and timbre for differences between conditions (p < 0.001). Furthermore, while intensity contrast was by itself not significant (p = .123), there was a significant difference between intensity and timbre for differences between conditions (p < 0.001). Taken together, these results explain the condition-by-feature interaction.

**Table 4.** Pairwise contrasts of P3a mean amplitudes between conditions for each feature separately, as well as pairwise comparisons between features for differences between conditions (contrasts of contrasts). Data is pooled from both groups. Significant contrasts are highlighted in bold and marked with an asterisk. Standard effect sizes (d) are calculated as the difference between conditions divided by the square root of the sum of the residual and the random effects variance.

| Feature | Contrast | Difference ($\mu V$) | CI 2.5% | CI 97.5% | t | d | p |
| --- | --- | --- | --- | --- | --- | --- | --- |
| Intensity | hi-hat - piano | -0.60 | -1.37 | 0.16 | -1.55 | -0.28 | 0.123 |
| Location | hi-hat - piano | -1.69 | -2.46 | -0.93 | -4.36 | -0.78 | **<0.001\*** |
| Timbre | hi-hat - piano | 2.94 | 2.18 | 3.71 | 7.58 | 1.35 | **<0.001\*** |
| (hi-hat - piano intensity) - (hi-hat - piano location) |  | 1.09 | -0.24 | 2.42 | 1.99 | 0.5 | 0.146 |
| (hi-hat - piano intensity) - (hi-hat - piano timbre) |  | -3.55 | -4.88 | -2.22 | -6.46 | -1.63 | **<0.001\*** |
| (hi-hat - piano location) - (hi-hat - piano timbre) |  | -4.64 | -5.97 | -3.31 | -8.44 | -2.13 | **<0.001\*** |

For mean P3a latencies (see Appendix 4), the main effects of condition and feature were significant, F(1,32) = 14.474, η^2^ = 0.033, p < .001 and F(2,64) = 45.605, η^2^ = 0.289, p < .001, respectively. No significant main effect was found for group F(1,32) = 0.297, p = .590. No significant interaction was found for group-by-feature F(2,64) = 1.203, p = .307. Interestingly, an interaction was found for group-by-condition F(1,32) = 5.094, η^2^ = 0.012, p = .031 and condition-by-feature F(2,64) = 4.892, η^2^ = 0.039, p = .011. No significant three-way interaction between group, condition and feature was observed, F(2,64) = 0.242, p = .786.

Post hoc pairwise comparisons (Table 5) revealed significant differences in latencies, contrasting hi-hat and piano conditions, for intensity with 31.59 milliseconds and location with 18.53 milliseconds (earlier P3a with piano tones in both cases). For timbre, no significant differences in latency between conditions were found (p = .440). Furthermore, while the difference between hi-hat and piano conditions was significant for the control group (23.12 milliseconds; 95% CI: 9.40-36.83; t(160) = 3.33; d = 0.57; p = .001), this was not the case for amusics (5.90 milliseconds; 95% CI: −7.81-19.62; t(160) = 0.85; d = 0.15; p = .396).

**Table 5.** Pairwise contrasts of P3a peak latencies between conditions for each feature separately, as well as pairwise comparisons between features for differences between conditions (contrasts of contrasts). Data is pooled from both groups. Significant contrasts are highlighted in bold and marked with an asterisk. Standard effect sizes (d) are calculated as the difference between conditions divided by the square root of the sum of the residual and the random effects variance.

| Feature | Contrast | Difference (ms) | CI 2.5% | CI 97.5% | t | d | p |
| --- | --- | --- | --- | --- | --- | --- | --- |
| Intensity | hi-hat - piano | 31.59 | 14.79 | 48.39 | 3.71 | 0.78 | **<.001\*** |
| Location | hi-hat - piano | 18.53 | 1.73 | 35.33 | 2.18 | 0.46 | **0.031\*** |
| Timbre | hi-hat - piano | -6.59 | -23.39 | 10.21 | -0.77 | -0.16 | 0.440 |
| (hi-hat - piano intensity) -<br>(hi-hat - piano location) |  | 13.06 | -16.04 | 42.16 | 1.09 | 0.32 | 0.838 |
| (hi-hat - piano intensity) -<br>(hi-hat - piano timbre) |  | 38.18 | 9.07 | 67.28 | 3.17 | 0.95 | **0.005\*** |
| (hi-hat - piano location) - (hi-hat<br>- piano timbre) |  | 25.12 | -3.99 | 54.22 | 2.09 | 0.62 | 0.115 |

## Discussion

In this study we found that the harmonicity of musical sounds enhances MMN in response to intensity, timbre and location deviants, both in control and amusic participants. This is consistent with the hypothesis that harmonicity influences the processing of auditory features other than pitch. In the following, an explanation based on precision-weighting of prediction errors will be discussed.

### Larger MMN for harmonic sounds

Harmonic sounds produced larger MMNs than inharmonic sounds, as evidenced by statistically significant main effects of condition (piano vs. hi-hat) on MMN amplitudes. Piano tones had clear harmonicity, whereas hi-hat tones had a more broadband spectrum (Figure 1). Other acoustic features were matched between the two: duration, impulsivity (sharp attack and fast decay as both types of sounds are generated by a single percussion), intensity, location; both had ecological validity. One possible interpretation for the larger MMN amplitudes in harmonic sounds comes from a predictive coding account of perception (Clark, 2015; Friston, 2005; Rao & Ballard, 1999). According to predictive coding, the brain employs a generative model to form predictions about the incoming sensory stimuli. The discrepancies between predictions and the sensory data, in the form of prediction errors, are used to update the generative model and enable better predictions in the future. This system is suggested to be hierarchical, with higher levels predicting the future states of lower levels, and the hierarchy is thought to be reflected in the neuroanatomical structure of the nervous system (Kanai et al., 2015). In predictive coding, the generative model is only updated by prediction errors with low self-estimated sensory uncertainty, or high *precision*. This *precision-weighting* process enables the system to “filter out” sensory data that is noisy, uncertain or otherwise unreliable and is analogous to attention (Kanai et al., 2015; Koelsch et al., 2019). ERP responses to deviances, such as MMN, are thought to be electrophysiological markers of ascending precision-weighted prediction errors (Garrido et al., 2007, 2007, 2009; Koelsch et al., 2019; Lecaignard et al., 2022; Winkler & Czigler, 2012(Garrido et al., 2007, 2008, 2009; Koelsch et al., 2019; Lecaignard et al., 2022; Winkler & Czigler, 2012; see also Grimm & Schroger, 2007 for a related, predictive account of MMN generation).

Precision of prediction errors is related to Shannon entropy, a metric that captures the amount of information content (uncertainty, expected surprise) of a given signal (Shannon, 1948). In individual sounds, entropy crucially depends on the amount of periodicity in the waveform shape and is thus directly related to harmonicity. Harmonic sounds have low spectral entropy because their waveforms can be reliably predicted using low amounts of information (as their frequency spectra consist of F0 and its integer multiples). Conversely, sounds that are inharmonic have higher spectral entropy and their waveforms are harder to predict (the frequency components do not form a harmonic series). Thus, the precision of prediction errors arising from deviances in inharmonic sounds is bound to be lower than in harmonic sounds. Consequently, the lower amplitudes of MMN signals for inharmonic sounds may result from a stronger down-weighting of sensory signals due to the spectral entropy of the acoustic input. Similar effects of information entropy on prediction error responses measured by MMN (or its magnetic counterpart, the MMNm) were reported previously for rhythmic (Lumaca et al., 2019) and melodic sequences (Quiroga-Martinez et al., 2019; Quiroga-Martinez et al., 2020). Furthermore, Takegata et al. (2008) investigated MMN in response to duration deviants for noise and other auditory stimuli. They found smaller MMN amplitudes in the noise condition in comparison to other (harmonic) stimuli. Although consistent with findings of the present study, these results were interpreted in terms of lower ecological validity of noise. The results of the present study suggest an effect of precision-weighting on prediction errors on a very short timescale during the unfolding of auditory signals, i.e. related to the instantaneous processing of the spectral content of sound.

### Differences between features in harmonic and non-harmonic sounds

We found that the strength of MMN responses was dependent on sound feature in different ways for the two types of sounds (piano vs. hi-hat), as evidenced by the significant interaction effect between condition and feature for MMN amplitudes and latencies. While the MMN was generally larger for harmonic than for inharmonic sounds, the differences between MMN amplitudes were less pronounced for timbre than for intensity and location deviants. Additionally, the MMN amplitudes for intensity and location deviants in the inharmonic condition were very small. MMN latencies were also dependent on sound feature, and this relationship was moderated by condition. For intensity deviants, harmonic sounds produced earlier peaks than inharmonic sounds, which is consistent with the idea that harmonic sounds are processed faster. Conversely, for timbre deviants, harmonic sounds produced later MMN peaks. This effect may be attributed to a large P3a response that followed the initial negativity (see below), possibly partly masking the MMN, especially for the hi-hat sound. No differences in latency between harmonic and inharmonic stimuli were found for the location deviants. As broadband noise stimuli are easier to locate than tones (Terhune, 1985), the broadband nature of the hi-hat sound may have counterbalanced the expected harmonicity effect. Additionally, MMN latency for location deviants was short (in comparison to latencies observed for classical MMN responses (Näätänen et al., 2007), suggesting that processing of location could be performed faster, on lower levels of the auditory hierarchy.

While MMN amplitudes were smaller for timbre than for other features, P3a amplitudes were larger, and were larger for hi-hat tones than piano tones. P3a indicates when the mental representation of a stimulus is updated (Donchin, 1981). Because of its longer latency, P3a is suggested to originate at higher levels of the auditory processing hierarchy than the MMN (Koelsch et al., 2019). The P3a is also associated with the initiation of an automatic attention-orienting response to stimulus deviance (Polich & Criado, 2006). Stronger P3a in our study could indicate a stronger implicit shift of attention to timbre deviants in comparison to other features. This shift may happen faster for timbre deviants, as evidenced by shorter P3a latencies. Furthermore, these strong and fast P3a responses for timbre could have interfered with activity in earlier latencies, potentially resulting in a diminished MMN response.

The distinct pattern of responses for timbre deviants may have to do with the filtering applied to the sounds to generate the deviants. Due to differences in the spectral characteristics of the piano and hi-hat, the filter changed these sounds in a different way. Because the hi-hat sound had a more uniformly distributed frequency spectrum, the filter introduced a timbral change that might have been perceived as a stronger deviance in comparison to the piano sound. It is thus possible that for the timbre condition, the different patterns of MMN and P3a might have resulted not from differences related to harmonicity, but from a more salient spectral change in the hi-hat condition. This problem is inherently related to comparing real-world harmonic and inharmonic sounds that (necessarily) differ in frequency spectra. One way to address this problem would be to investigate natural sounds with artificially manipulated harmonic components (as in Popham, 2018). Another approach could involve fully synthesized sounds with carefully controlled manipulations in the frequency spectra, yet this may negatively impact the ecological validity of the stimuli. This filter-related restriction however applies only to timbre deviants and does not extend to other auditory features.

### Preserved MMN and P3a in congenital amusia for harmonic and non-harmonic sounds

We found significant MMN and P3a responses, both in CA and control participants and no significant main effects of group or group-condition, group-feature or group-feature-condition interactions. This was true for mean amplitudes as well as mean latencies in both MMN and P3a analyses, with the exception of a group-by-condition interaction in P3a latency (discussed below). Whether or not the processing deficits in congenital amusia extend to the processing of harmonicity is debated. It was reported that amusics are insensitive to harmonicity cues, contrarily to beating cues (Cousineau et al., 2012; but see Graves et al., 2021), and they might rely on roughness cues more than harmonicity cues to evaluate the pleasantness of a sound (Marin et al., 2012). On the other hand, the identification of pitch processing areas with fMRI by contrasting harmonic sounds with noises lead to results very similar to that obtained in controls (Norman-Haignere et al., 2016), yet with subtle differences revealed by multivariate analyses (Albouy, Caclin, et al., 2019). During active pitch short-term memory tasks, functional impairments are observed in a fronto-temporal network in congenital amusia (Albouy et al., 2013; Albouy, Peretz, et al., 2019). Our results suggest that individuals with CA do not differ from matched controls in terms of early neural processing of auditory features other than pitch (see Quiroga-Martinez et al., 2021, for converging evidence). Event-related potential studies with pitch deviants in classical oddball paradigms also report intact MMN responses in CA listeners, indicating normal pitch processing at an early level of auditory processing (Moreau et al., 2013; Zendel et al., 2015), whereas abnormalities might be observed in more complex sound sequences (Quiroga-Martinez et al., 2021). Taken together, these results provide further evidence that deficits in CA might mainly result from abnormalities at higher levels of the perceptual hierarchy, possibly related to impaired top-down connectivity in the frontotemporal network (Albouy et al., 2013; Peretz, 2016). Our results can be also interpreted as evidence that low-level auditory processing in CA remains greatly spared, not only for pitch, but also for harmonicity, even though the deficit might lead to delayed processing with impaired auditory cortex contributions (see Albouy et al., 2013). This is consistent with the hypothesis that CA individuals might not have conscious access to the results of low-level processing and might thus be less able to use them in perceptual decision making (Peretz, 2016).

In P3a latency analysis, we found a significant group-by-condition interaction. This was accompanied with a significant main effect of condition (piano sounds yielded generally shorter P3a latencies than hi-hat sounds) but not a main effect of group. The strength of this effect is however very small (η^2^ = 0.012), much smaller than the main effects of condition (η^2^ = 0.033) or feature (η^2^ = 0.289). This interaction can be interpreted as suggesting that for inharmonic sounds, the CA individuals require more processing time to consciously detect deviance. However, because of the small effect size further research is needed to clarify this finding.

### Limitations and conclusions

A limitation of our study is that subjects listened to the stimuli passively, without any perceptual tasks. Our goal was to focus on low-level processing of auditory deviances and attention-orientation as measured with the MMN and P3a, respectively. We chose a passive listening approach because it is optimal for evaluating the MMN. More active tasks (such as actively detecting deviants with button-presses) could be used to investigate neural processing that involves higher levels of the perceptual hierarchy, with a focus on later ERPs. These types of tasks could also arguably reveal differences in processing specific to CA that were not found in the current study. Finally, it would be beneficial to investigate a more diverse set of auditory stimuli with different levels of inharmonicity to better understand the relationship between the spectral properties of sound and the MMN. Piano and hi-hat sounds differ not only in harmonicity, but in other features as well (e.g., amplitude envelope, spectral content). Future studies may attempt to control for this issue using either complex tones with jittered frequencies, or use natural sounds with artificially manipulated harmonics. In a recent study with synthetic tones (Graves et al., in revision), we observed larger MMNs for harmonic deviants within a sequence of inharmonic standards than the reverse (inharmonic deviants within a sequence of harmonic standards), paving the way for future studies were individual sound entropies could be carefully controlled in oddball sequences.

The results of the present study provide neurophysiological evidence consistent with the notion that harmonicity plays an important role in human auditory perception, starting in early stages of sound processing in the auditory networks. A detailed investigation into the relationships between early auditory ERPs and harmonicity can provide valuable information about the nature of auditory information processing in the brain.

## Abbreviations

ANOVA: analysis of variance
CA: congenital amusia
CI: confidence interval
Df: degrees of freedom
EEG: electroencephalography
ERP: event related potential
F0: fundamental frequency
ICA: independent component analysis
M: mean
MBEA: Montreal Battery of Evaluation of Amusia
MMN: mismatch negativity
PDT: pitch discrimination threshold
SD: standard deviation

## CrediT authorship contribution statement

David R. Quiroga-Martinez: Conceptualization, Methodology, Software, Formal analysis, Data curation, Writing - original draft, Visualization, Investigation. Krzysztof Basiński: Software, Writing - original draft, Visualization. Jonathan Nasielski: Software, Writing - original draft, Visualization. Barbara Tillmann: Conceptualization, Supervision, Writing - original draft. Elvira Brattico: Conceptualization, Supervision, Writing - original draft. Fanny Cholvy - Investigation, Data curation. Lesly Fornoni - Investigation, Data curation. Peter Vuust: Conceptualization, Methodology, Supervision, Writing - original draft. Anne Caclin: Conceptualization, Methodology, Writing - original draft, Supervision.

## Open practices

The code and materials employed to conduct the experiment and analyses presented here can be found at: http://doi.org/10.17605/OSF.IO/JE4XZ. Due to data protection regulations, data cannot be publicly shared, but can be made privately available upon reasonable request.

## Appendices

**Appendix 1.** ANOVA table for MMN amplitude analysis: main effects, two-way and three-way interaction effects on group, condition, and feature. Significant effects are marked with an asterisk.

| Effect | DFn | DFd | SSn | SSd | F | p | ges |
| --- | --- | --- | --- | --- | --- | --- | --- |
| (Intercept) | 1 | 32 | 1098.317 | 278.878 | 126.027 | <.001* | 0.687 |
| group | 1 | 32 | 0.55 | 278.878 | 0.063 | .803 | 0.001 |
| condition | 1 | 32 | 256.001 | 81.006 | 101.129 | <.001* | 0.338 |
| feature | 2 | 64 | 3.318 | 93.889 | 1.131 | .329 | 0.007 |
| group:condition | 1 | 32 | 1.231 | 81.006 | 0.486 | .491 | 0.002 |
| group:feature | 2 | 64 | 7.513 | 93.889 | 2.561 | .085 | 0.015 |
| condition:feature | 2 | 64 | 36.497 | 47.698 | 24.485 | <.001* | 0.068 |
| group:condition:feature | 2 | 64 | 1.155 | 47.698 | 0.775 | .465 | 0.002 |

**Appendix 2.** ANOVA table for MMN latency analysis: main effects, two-way and three-way interaction effects on group, condition, and feature. Significant effects are marked with an asterisk.

| Effect | DFn | DFd | SSn | SSd | F | p | ges |
| --- | --- | --- | --- | --- | --- | --- | --- |
| (Intercept) | 1 | 32 | 2869123.77 | 33201.059 | 2765.332 | <.001* | 0.955 |
| group | 1 | 32 | 202.005 | 33201.059 | 0.195 | 0.662 | 0.001 |
| condition | 1 | 32 | 313.77 | 21153.333 | 0.475 | 0.496 | 0.002 |
| feature | 2 | 64 | 10732.392 | 37121.471 | 9.252 | <.001* | 0.073 |
| group:condition | 1 | 32 | 42.397 | 21153.333 | 0.064 | 0.802 | 0.0 |
| group:feature | 2 | 64 | 291.804 | 37121.471 | 0.252 | 0.778 | 0.002 |
| condition:feature | 2 | 64 | 15958.745 | 44329.667 | 11.52 | <.001* | 0.105 |
| group:condition:feature | 2 | 64 | 632.588 | 44329.667 | 0.457 | 0.635 | 0.005 |

**Appendix 3.** ANOVA table for P3a amplitude analysis: main effects, two-way and three-way interaction effects on group, condition, and feature. Significant effects are marked with an asterisk.

| Effect | DFn | DFd | SSn | SSd | F | p | ges |
| --- | --- | --- | --- | --- | --- | --- | --- |
| (Intercept) | 1 | 32 | 1338.018 | 502.816 | 85.154 | **<.001\*** | 0.594 |
| group | 1 | 32 | 3.505 | 502.816 | 0.223 | 0.64 | 0.004 |
| condition | 1 | 32 | 2.378 | 88.18 | 0.863 | 0.36 | 0.003 |
| feature | 2 | 64 | 136.148 | 177.249 | 24.58 | **<.001\*** | 0.13 |
| group:condition | 1 | 32 | 0.94 | 88.18 | 0.341 | 0.563 | 0.001 |
| group:feature | 2 | 64 | 2.87 | 177.249 | 0.518 | 0.598 | 0.003 |
| condition:feature | 2 | 64 | 199.872 | 144.797 | 44.171 | **<.001\*** | 0.18 |
| group:condition:feature | 2 | 64 | 6.755 | 144.797 | 1.493 | 0.232 | 0.007 |

**Appendix 4.**
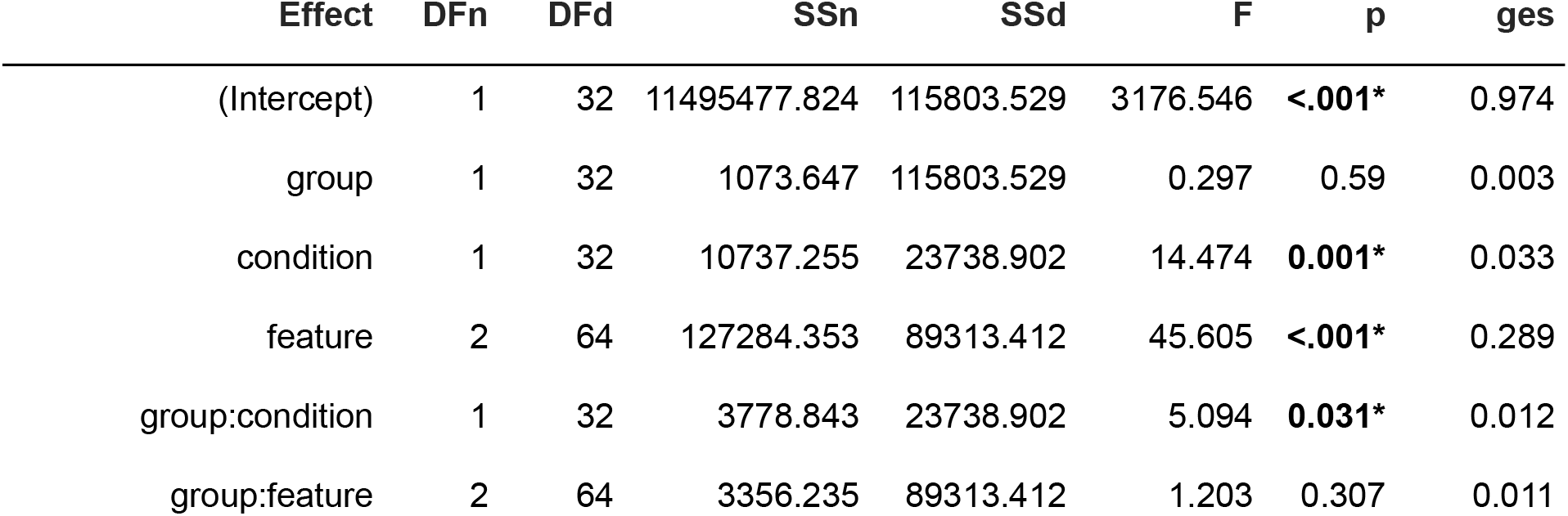
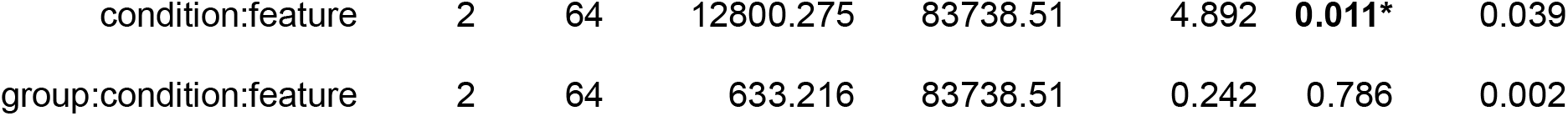
ANOVA table for P3a latency analysis: main effects, two-way and three-way interaction effects on group, condition, and feature. Significant effects are marked with an asterisk.

| Effect | DFn | DFd | SSn | SSd | F | p | ges |
| --- | --- | --- | --- | --- | --- | --- | --- |
| (Intercept) | 1 | 32 | 11495477.824 | 115803.529 | 3176.546 | **<.001\*** | 0.974 |
| group | 1 | 32 | 1073.647 | 115803.529 | 0.297 | 0.59 | 0.003 |
| condition | 1 | 32 | 10737.255 | 23738.902 | 14.474 | **0.001\*** | 0.033 |
| feature | 2 | 64 | 127284.353 | 89313.412 | 45.605 | **<.001\*** | 0.289 |
| group:condition | 1 | 32 | 3778.843 | 23738.902 | 5.094 | **0.031\*** | 0.012 |
| group:feature | 2 | 64 | 3356.235 | 89313.412 | 1.203 | 0.307 | 0.011 |
| condition:feature | 2 | 64 | 12800.275 | 83738.51 | 4.892 | **0.011\*** | 0.039 |
| group:condition:feature | 2 | 64 | 633.216 | 83738.51 | 0.242 | 0.786 | 0.002 |

## References

Albouy, P., Caclin, A., Norman-Haignere, S. V., Lévêque, Y., Peretz, I., Tillmann, B., & Zatorre, R. J. (2019). Decoding Task-Related Functional Brain Imaging Data to Identify Developmental Disorders: The Case of Congenital Amusia. Frontiers in Neuroscience, 13, 1165. https://doi.org/10.3389/fnins.2019.01165

Albouy, P., Mattout, J., Bouet, R., Maby, E., Sanchez, G., Aguera, P.-E., Daligault, S., Delpuech, C., Bertrand, O., Caclin, A., & Tillmann, B. (2013). Impaired pitch perception and memory in congenital amusia: The deficit starts in the auditory cortex. Brain, 136(5), 1639–1661. https://doi.org/10.1093/brain/awt082

Albouy, P., Peretz, I., Bermudez, P., Zatorre, R. J., Tillmann, B., & Caclin, A. (2019). Specialized neural dynamics for verbal and tonal memory: FMRI evidence in congenital amusia. Human Brain Mapping, 40(3), 855–867.

Bates, D., Mächler, M., Bolker, B., & Walker, S. (2015). Fitting Linear Mixed-Effects Models Using lme4. Journal of Statistical Software, 67(1), 1–48. https://doi.org/10.18637/jss.v067.i01

Bregman, A. S. (1994). Auditory Scene Analysis: The Perceptual Organization of Sound. MIT Press.

Clark, A. (2015). Surfing uncertainty: Prediction, action, and the embodied mind. Oxford University Press.

Cousineau, M., McDermott, J. H., & Peretz, I. (2012). The basis of musical consonance as revealed by congenital amusia. Proceedings of the National Academy of Sciences, 109(48), 19858–19863. https://doi.org/10.1073/pnas.1207989109

Donchin, E. (1981). Surprise! … Surprise? Psychophysiology, 18(5), 493–513. https://doi.org/10.1111/j.1469-8986.1981.tb01815.x

Friston, K. J. (2005). A theory of cortical responses. Philosophical Transactions of the Royal Society B: Biological Sciences, 360(1456), 815–836. https://doi.org/10.1098/rstb.2005.1622

Garrido, M. I., Friston, K. J., Kiebel, S. J., Stephan, K. E., Baldeweg, T., & Kilner, J. M. (2008). The functional anatomy of the MMN: A DCM study of the roving paradigm. NeuroImage, 42(2), 936–944. https://doi.org/10.1016/j.neuroimage.2008.05.018

Garrido, M. I., Kilner, J. M., Kiebel, S. J., & Friston, K. J. (2007). Evoked brain responses are generated by feedback loops. Proceedings of the National Academy of Sciences, 104(52), 20961–20966. https://doi.org/10.1073/pnas.0706274105

Garrido, M. I., Kilner, J. M., Stephan, K. E., & Friston, K. J. (2009). The mismatch negativity: A review of underlying mechanisms. Clinical Neurophysiology, 11.

Gramfort, A., Luessi, M., Larson, E., Engemann, D. A., Strohmeier, D., Brodbeck, C., Parkkonen, L., & Hämäläinen, M. S. (2014). MNE software for processing MEG and EEG data. NeuroImage, 86, 446–460. https://doi.org/10.1016/j.neuroimage.2013.10.027

Graves, J. E., Pralus, A., Fornoni, L., Oxenham, A. J., Tillmann, B., & Caclin, A. (2021). Consonance perception in congenital amusia: Behavioral and brain responses to harmonicity and beating cues [Preprint]. Neuroscience. https://doi.org/10.1101/2021.11.15.468620

Grimm, S., & Schroger, E. (2007). The processing of frequency deviations within sounds: Evidence for the predictive nature of the Mismatch Negativity (MMN) system. Restorative Neurology and Neuroscience, 25, 241–249.

Huron, D. (2016). Voice Leading: The Science behind a Musical Art. MIT Press.

Kanai, R., Komura, Y., Shipp, S., & Friston, K. (2015). Cerebral hierarchies: Predictive processing, precision and the pulvinar. Philosophical Transactions of the Royal Society B: Biological Sciences, 370(1668), 20140169. https://doi.org/10.1098/rstb.2014.0169

Kliuchko, M., Heinonen-Guzejev, M., Vuust, P., Tervaniemi, M., & Brattico, E. (2016). A window into the brain mechanisms associated with noise sensitivity. Scientific Reports, 6(1), 39236. https://doi.org/10.1038/srep39236

Koelsch, S., Vuust, P., & Friston, K. (2019). Predictive Processes and the Peculiar Case of Music. Trends in Cognitive Sciences, 23(1), 63–77. https://doi.org/10.1016/j.tics.2018.10.006

Lawrence, M. A. (2016). ez: Easy Analysis and Visualization of Factorial Experiments (4.4-0) [Computer software]. https://CRAN.R-project.org/package=ez

Lecaignard, F., Bertrand, O., Caclin, A., & Mattout, J. (2022). Neurocomputational Underpinnings of Expected Surprise. The Journal of Neuroscience, 42(3), 474–486. https://doi.org/10.1523/JNEUROSCI.0601-21.2021

Lenth, R. V., Buerkner, P., Herve, M., Love, J., Riebl, H., & Singmann, H. (2021). emmeans: Estimated Marginal Means, aka Least-Squares Means (1.5.5-1) [Computer software]. https://CRAN.R-project.org/package=emmeans

Lumaca, M., Trusbak Haumann, N., Brattico, E., Grube, M., & Vuust, P. (2019). Weighting of neural prediction error by rhythmic complexity: A predictive coding account using mismatch negativity. European Journal of Neuroscience, 49(12), 1597–1609. https://doi.org/10.1111/ejn.14329

Marin, M. M., Gingras, B., & Stewart, L. (2012). Perception of musical timbre in congenital amusia: Categorization, discrimination and short-term memory. Neuropsychologia, 50(3), 367–378. https://doi.org/10.1016/j.neuropsychologia.2011.12.006

McDermott, J. H. (2018). Audition. In J. T. Wixted (Ed.), Stevens’ Handbook of Experimental Psychology and Cognitive Neuroscience (pp. 1–57). John Wiley & Sons, Inc. https://doi.org/10.1002/9781119170174.epcn202

McPherson, M. J., Grace, R. C., & McDermott, J. H. (2020). Harmonicity aids hearing in noise [Preprint]. Neuroscience. https://doi.org/10.1101/2020.09.30.321000

McPherson, M. J., & McDermott, J. H. (2020a). Time-dependent discrimination advantages for harmonic sounds suggest efficient coding for memory. Proceedings of the National Academy of Sciences, 117(50), 32169–32180. https://doi.org/10.1073/pnas.2008956117

McPherson, M. J., & McDermott, J. H. (2020b). Time-dependent discrimination advantages for harmonic sounds suggest efficient coding for memory. Proceedings of the National Academy of Sciences, 117(50), 32169–32180. https://doi.org/10.1073/pnas.2008956117

Moreau, P., Jolicœur, P., & Peretz, I. (2013). Pitch discrimination without awareness in congenital amusia: Evidence from event-related potentials. Brain and Cognition, 81(3), 337–344. https://doi.org/10.1016/j.bandc.2013.01.004

Näätänen, R., Paavilainen, P., Rinne, T., & Alho, K. (2007). The mismatch negativity (MMN) in basic research of central auditory processing: A review. Clinical Neurophysiology, 118(12), 2544–2590. https://doi.org/10.1016/j.clinph.2007.04.026

Näätänen, R., Pakarinen, S., Rinne, T., & Takegata, R. (2004). The mismatch negativity (MMN): Towards the optimal paradigm. Clinical Neurophysiology, 115(1), 140–144. https://doi.org/10.1016/j.clinph.2003.04.001

Norman-Haignere, S. V., Albouy, P., Caclin, A., McDermott, J. H., Kanwisher, N. G., & Tillmann, B. (2016). Pitch-Responsive Cortical Regions in Congenital Amusia. Journal of Neuroscience, 36(10), 2986–2994. https://doi.org/10.1523/JNEUROSCI.2705-15.2016

Oxenham, A. J. (2012). Pitch Perception. Journal of Neuroscience, 32(39), 13335–13338. https://doi.org/10.1523/JNEUROSCI.3815-12.2012

Paavilainen, P. (2013). The mismatch-negativity (MMN) component of the auditory event-related potential to violations of abstract regularities: A review. International Journal of Psychophysiology, 88(2), 109–123. https://doi.org/10.1016/j.ijpsycho.2013.03.015

Peretz, I. (2016). Neurobiology of Congenital Amusia. Trends in Cognitive Sciences, 20(11), 857–867. https://doi.org/10.1016/j.tics.2016.09.002

Peretz, I., Champod, A. S., & Hyde, K. (2003). Varieties of Musical Disorders. Annals of the New York Academy of Sciences, 999(1), 58–75. https://doi.org/10.1196/annals.1284.006

Polich, J., & Criado, J. R. (2006). Neuropsychology and neuropharmacology of P3a and P3b. International Journal of Psychophysiology, 60(2), 172–185. https://doi.org/10.1016/j.ijpsycho.2005.12.012

Popham, S. (2018). Inharmonic speech reveals the role of harmonicity in the cocktail party problem. NATURE COMMUNICATIONS, 15.

Quiroga Martinez, D. R., C. Hansen N., Højlund, A., Pearce, M., Brattico, E., & Vuust, P. (2020). Musical prediction error responses similarly reduced by predictive uncertainty in musicians and non musicians. European Journal of Neuroscience, 51(11), 2250–2269. https://doi.org/10.1111/ejn.14667

Quiroga-Martinez, D. R., Hansen, N. C., Højlund, A., Pearce, M. T., Brattico, E., & Vuust, P. (2019). Reduced prediction error responses in high-as compared to low-uncertainty musical contexts. Cortex, 120, 181–200. https://doi.org/10.1016/j.cortex.2019.06.010

Quiroga-Martinez, D. R., Tillmann, B., Brattico, E., Cholvy, F., Fornoni, L., Vuust, P., & Caclin, A. (2021). Listeners with congenital amusia are sensitive to context uncertainty in melodic sequences. Neuropsychologia, 158, 107911. https://doi.org/10.1016/j.neuropsychologia.2021.107911

R Core Team. (2021). R: The R Project for Statistical Computing. https://www.r-project.org/

Rao, R. P. N., & Ballard, D. H. (1999). Predictive coding in the visual cortex: A functional interpretation of some extra-classical receptive-field effects. Nature Neuroscience, 2(1), 79–87. https://doi.org/10.1038/4580

Shannon, C. E. (1948). A Mathematical Theory of Communication. 55.

Terhune, J. M. (1985). Localization of Pure Tones and Click Trains by Untrained Humans. Scandinavian Audiology, 14(3), 125–131. https://doi.org/10.3109/01050398509045933

Tillmann, B., Schulze, K., & Foxton, J. (2009). Congenital amusia: A short-term memory deficit for non-verbal, but not verbal sounds. Brain and Cognition. https://doi.org/10.1016/j.bandc.2009.08.003

Vallat, R. (2021). AntroPy: Entropy and complexity of (EEG) time-series in Python (0.1.4) [Computer software]. https://raphaelvallat.com/antropy/

Vuust, P., Brattico, E., Glerean, E., Seppänen, M., Pakarinen, S., Tervaniemi, M., & Näätänen, R. (2011). New fast mismatch negativity paradigm for determining the neural prerequisites for musical ability. Cortex, 47(9), 1091–1098. https://doi.org/10.1016/j.cortex.2011.04.026

Wightman, F. L., & Green, D. M. (1974). The Perception of Pitch: The pitch of a sound wave is closely related to its frequency or periodicity—but the exact nature of that relation remains a mystery. American Scientist, 62(2), 208–215.

Winkler, I., & Czigler, I. (2012). Evidence from auditory and visual event-related potential (ERP) studies of deviance detection (MMN and vMMN) linking predictive coding theories and perceptual object representations. International Journal of Psychophysiology, 83(2), 132–143. https://doi.org/10.1016/j.ijpsycho.2011.10.001

Zendel, B. R., Lagrois, M.-E., Robitaille, N., & Peretz, I. (2015). Attending to Pitch Information Inhibits Processing of Pitch Information: The Curious Case of Amusia. Journal of Neuroscience, 35(9), 3815–3824. https://doi.org/10.1523/JNEUROSCI.3766-14.2015

